# Repertoire-based mapping and time-tracking of helper T cell subsets in scRNA-Seq

**DOI:** 10.1101/2023.10.24.563704

**Authors:** D.K. Lukyanov, V.V. Kriukova, K. Ladell, I.A. Shagina, D.B. Staroverov, B.E. Minasian, A.S. Fedosova, P. Shelyakin, O.N. Suchalko, A.Y. Komkov, K.A. Blagodatskikh, K.L. Miners, O.V. Britanova, A. Franke, D.A. Price, D.M. Chudakov

## Abstract

The functional programs selected by CD4+ helper (Th) T cell clones fundamentally determine the architecture of the immune response to distinct challenges. Advances in scRNA-Seq have enhanced our understanding of the diversity of these programs, yet the correspondence between scRNA-Seq clusters and previously characterized Th subsets remains unclear. In this study, we use immune repertoires to position phenotypically sorted Th subsets within scRNA-Seq data from three healthy donors. This approach, termed TCR-Track, and accurately maps Th1, Th1-17, Th17, Th22, Th2a, Th2, Tfh, and Treg subsets, outperforming CITE-Seq-based mapping. Remarkably, the mapping is tightly focused on specific scRNA-Seq clusters despite a four-year interval between the sorting of subsets and the effector CD4+ scRNA-Seq experiment. Thus, while transient T cell plasticity is commonly observed in functionally active T cell populations, TCR-Track reveals high intrinsic program sustainability of Th clones circulating in peripheral blood. Repertoire overlap analysis at the scRNA-Seq level confirms that circulating Th1, Th2, Th2a, Th17, Th22, and Treg subsets are clonally independent. However, a prominent clonal overlap between corresponding clusters indicates that cytotoxic CD4+ T cells differentiate from Th1 clones. More specifically, we demonstrate that sorted CCR10+ Th cells correspond to a specific Th22 scRNA-Seq cluster, while CCR10-CCR6+CXCR3-CCR4+ cells, traditionally sorted as the Th17 subset, represent a mixture of *bona fide* Th17 and clonally unrelated CCR10^low^ Th22 cells, which may have confounded investigators in previous studies. This clear distinction of Th17 and Th22 subsets should influence vaccine and T cell based therapies development. Additionally, we show that SARS-CoV-2 infection is associated with transient IFN type 1 activation of naive CD4+ T cells, and an increased proportion of effector IFN-induced Th cells is associated with a moderate course of the disease but remains low in critical COVID-19 cases. Using integrated scRNA-Seq, TCR-Track, and CITE-Seq data from 122 donors, we provide a comprehensive Th scRNA-Seq reference that should facilitate further investigation of Th subsets in fundamental and clinical studies.

## Introduction

Clonal populations of CD4^+^ T helper (Th) cells orchestrate the course of an immune response via specific interactions with peptide epitopes presented in complex with major histocompatibility class II (MHCII) molecules. Their functional and antigen-specific diversity allows them to guide both classical (B cells, dendritic cells, macrophages) and non-classical (endothelial cells, epithelial cells, granulocytes) antigen-presenting cells to optimize effector functionality^1, 2, 3^. Inappropriate Th responses to certain antigens have been associated with impaired pathogen clearance^4, 5, 6^, inefficient response to vaccination^7^, acute and chronic hypersensitivity and inflammation^8, 9, 10^, inflammaging^11^, autoimmunity^12, 13, 14, 15^, and cancer^16^. Accordingly, when investigating T cell responses, it is critical not only to quantify the magnitude of the antigen-specific T cell clonal expansion but also to understand functional programs and related phenotypes of the responding and memory helper T cells.

Single-cell RNA-Seq (scRNA-Seq) techniques are shedding light on the diversity of Th cell programs^17, 18, 19, 20^, among which classical subsets previously described on the basis of cytokine release profiles and patterns of surface marker expression have to find their place. The expression of transcripts encoding characteristic surface markers is often low in scRNA-Seq datasets, with indirect correlations between mRNA abundance and protein density^21, 22^. This problem can be overcome to some extent by the incorporation of protein-level expression data into single-cell experiments, a task that is essentially implemented in CITE-Seq technology, which makes use of barcoded antibodies directed against markers of interest expressed on the cell surface^23, 24, 25^.

Here, we report an approach to the mapping and clonal tracking of sorted lymphocyte subsets within scRNA-Seq data, termed TCR-Track. This approach makes use of immune repertoires from sorted lymphocyte subsets, which are then mapped to scRNA-Seq+TCR-Seq datasets obtained from the same donors, employing natural barcodes in the form of sequence-defined TCRs. We demonstrate that this method accurately maps phenotypically defined Th subsets within scRNA-Seq landscape. We further integrate scRNA-Seq, TCR-Track, and CITE-Seq outputs of 122 donors to provide a comprehensive Th scRNA-Seq reference dataset.

TCR-Track also allowed us to trace the positioning of subset-specific T cell clones obtained from the deep bulk TCR profiling of sorted subsets after a 4 year interval. The latter approach, complementing concepts describing plasticity of tissue resident T cells^26, 27^, reveals surprisingly high long-term program stability of CD4+ T cell clones circulating in human peripheral blood.

In general, based on the integrated reference, we expand upon previous observations of the clonal independence of circulating Th1, Th2, Th2a, Th17, Th22, and Treg subsets^28^. We also disentangle interrelations between the subsets classically sorted as Th17 and Th22, and demonstrate a prominent clonal overlap between Th1 and cytotoxic Th subsets, supporting their common lineage^29, 30^.

Finally, we report an association between SARS-CoV-2 infection and transient IFN type 1 activation of naive CD4+ T cells, as well as a link between an increased proportion of IFN-induced Th cells and a moderate course of the disease.

## Results

### TCR-Track

Functional subtypes of human helper T cells are classically distinguished based on the surface markers, such as CD127, CD25, CCR10, CXCR5, CXCR3, CCR6, CCR4, and CRTh2. Here we conceptualized the annotation of Th cells in scRNA-Seq data powered by overlapping scTCR-Seq repertoires and TCR repertoires of FACS-sorted Th cell subsets, TCR-Track. We hypothesized that, due to relatively high phenotypic stability of helper T cell subsets^28^, this approach could map sorted Th subsets within scRNA-Seq data (**Figure 1**).

**Figure 1.**
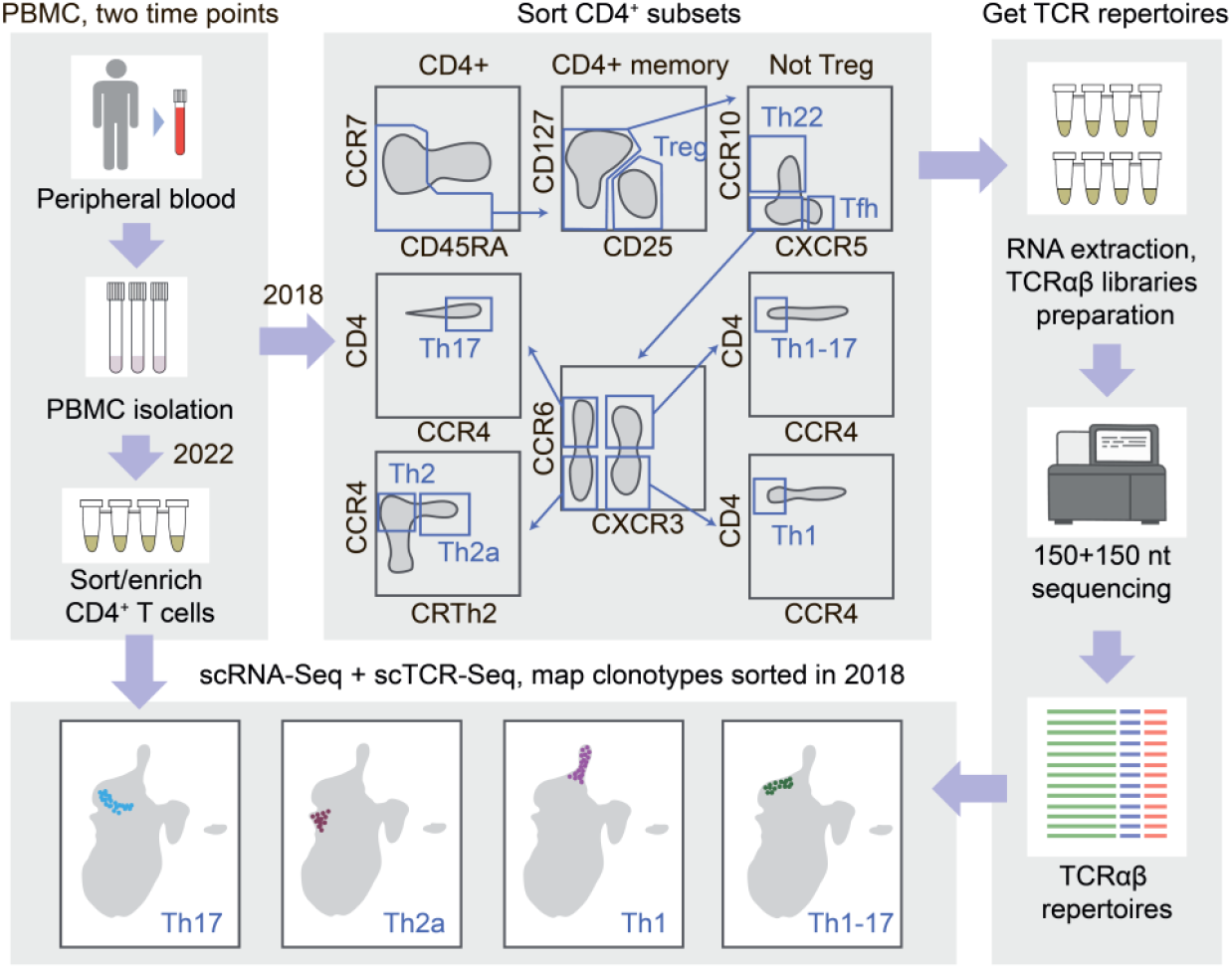
Scheme of the TCR-based mapping of sorted Th subsets in scRNA-Seq data.

We exploited previously obtained TCRα and TCRβ repertoires of Th1, Th17, Th1-17, Th22, Th2, Th2a, Treg, and Tfh CD4+ T cell subsets thoroughly sorted from peripheral blood of healthy donors^28^. Three participants of the latter study were available for repeated blood donation at the moment of the current study. We performed paired scRNA-Seq and scTCR-Seq profiling from their sorted effector/memory CD4+ T cells (gated as CD4+, NOT CCR7+CD45RA+ cells to deplete naive T cells). The obtained scRNA-Seq data has then been integrated with the CD4+ scRNA-Seq reported in Ref. 31 in order to: 1) increase the power of the downstream analysis such as clustering and UMAP visualization on a larger number of donors and 2) enable comparison/complementing of mapping with the CITE-Seq method employed in the latter work. This resulted in a reference dataset composed of 147,677 cells (**Figure 2A**), without notable donor-specific or study-specific batch effects (**Figures S1A**).

**Figure 2.**
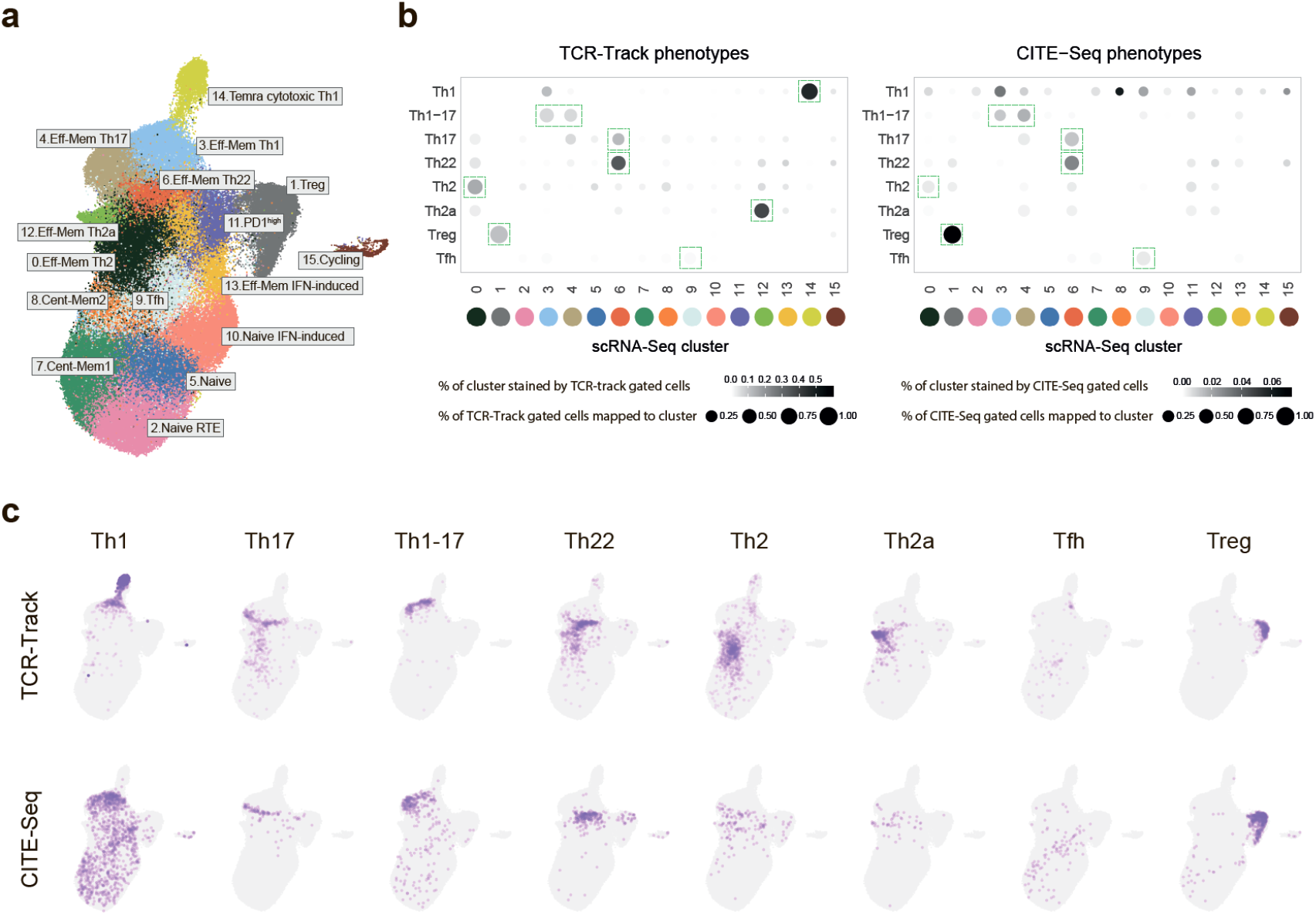
Mapping the classic Th subsets with scRNA-Seq. **a.** UMAP visualization of the reference scRNA-Seq dataset of peripheral blood Th cells. Dataset built via Seurat integration of publicly available and our scRNA-Seq data. The proposed classification is based on previous knowledge and findings of the current work. **b**. Dot plots summarizing the positioning of sorted (TCR-Track) and *in silico* gated (CITE-Seq) Th subsets within the scRNA-Seq clusters shown on the panel (c). For normalization, we used 20,000 randomly selected scRNA-Seq cells with matched CITE-Seq or TCR-Track data for each plot. Dot intensity shows the stained proportion of the scRNA-Seq cluster. Dot size shows the proportion of TCR-Track-identified or proportion of *in silico* CITE-Seq-based gated scRNA-Seq cells mapped to the scRNA-Seq cluster. Green dashed rectangles indicate dominating scRNA-Seq cluster. **c**. UMAP plots showing the localization of TCR-Track and CITE-Seq defined subsets. TCRβ clonotypes were used to define Th subsets in TCR-Track method. Expression of the surface markers was used to gate Th subsets in the CITE-Seq-based annotation. The color intensity in TCR-Track is proportional to the clonal frequencies in the original sorted Th bulk TCRβ repertoires.

In order to link scRNA-Seq clusters to the classic Th phenotypes, we mapped TCR clonotypes of the sorted Th subsets using scTCRs as natural barcodes. Remarkably, T cell clones from each of the sorted Th subpopulations formed clearly defined spots on the scRNA-Seq UMAP (**Figure 2B-D, Figures S1B,C**). This allowed us to exploit TCR-Track to build “correspondence between the nomenclatures” of classic immunology (based on the surface molecules and flow cytometry; these are the subsets that can be physically sorted and investigated *in vitro*) and scRNA-Seq landscape (based on gene expression at the RNA level), filling the gaps of potential miscorrelation between the surface proteins, FACS sorting sensitivity, and mRNA expression levels.

Most of the scRNA-Seq clusters showed low clonality in the scTCR-Seq data, thus excluding strong clonal-driven biases in the TCR-Track-based annotation (**Figure 3A**).

**Figure 3.**
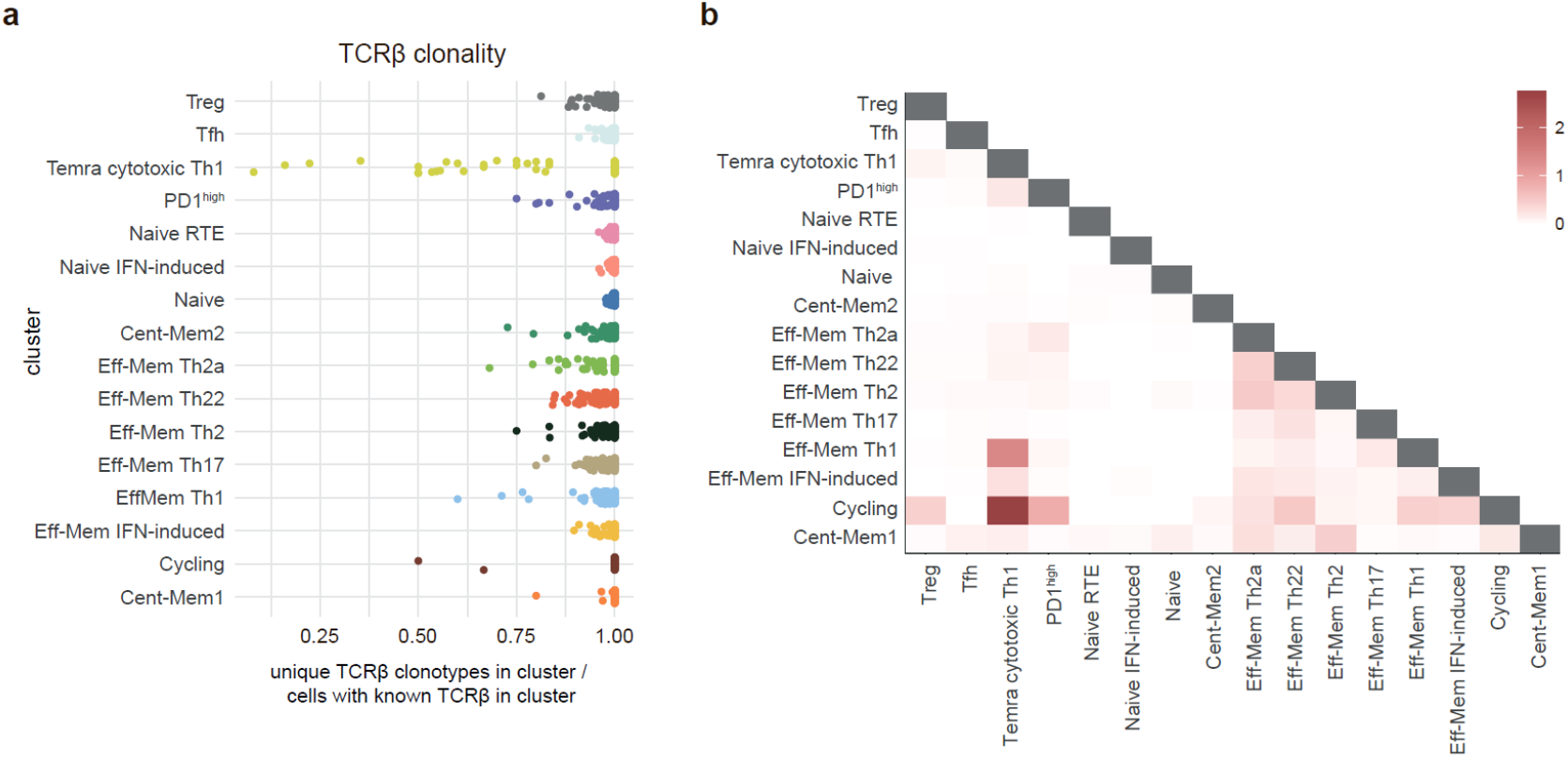
Clonality and overlap between Th clusters. **a.** Relative clonality of scRNA-Seq Th clusters represented as unique TCRβ CDR3/cell count ratio. Each dot represents one donor. **b**. Heatmap visualization of clonal (nucleotide-defined TCRβ CDR3) overlaps between scRNA-Seq clusters measured as the number of shared clonotypes between the clusters divided by the number of clonotypes in each cluster (D metrics in VDJtools). The D metric values are log_2_(1+x) transformed^40, 41^.

Additionally, we annotated the same surface-marker defined Th subsets within scRNA-Seq using CITE-Seq. For this, we used CITE-Seq data from Ref. 31, and performed sequential *in silico* sorting-like gating as shown in **Figure S2**. TCR-Track was generally consistent with the CITE-Seq data analysis, but resulted in much more accurate cluster annotations (**Figure 2B,C, Figures S1B-D, S3, S4**). This suggests that TCR-Track represents more reliable method of T cell subsets mapping, with many additional options provided by opportunity to track T cell clones in time and space and across different methods, employing TCR as a natural barcode.

Based on the integrated data of scRNA-Seq, TCR-Track, and CITE-Seq outputs of 122 donors, taking into account findings and considerations of Sakaguchi and colleagues^18^, we propose a comprehensive peripheral blood Th scRNA-Seq reference map with 16 major Th clusters (**Figure 2A, Supplementary Table 1, Figures S4, S5**), as described in more detail below.

### Correspondence between classic Th subsets and scRNA-Seq clusters

TCR-Track positions TCRβ clonotypes of sorted Treg (CD25^high^CD127^low^), Th22 (CXCR5-CCR10+), Th2 (CCR10-CXCR5-CCR6-CXCR3-CCR4+CRTh2-), and Th2a (CCR10-CXCR5-CCR6-CXCR3-CCR4+

CRTh2+) cells to corresponding unique scRNA-Seq clusters, clearly fixing their localization in the peripheral blood scRNA-Seq landscape (**Figure 2**).

Sorted Th1 (CCR10-CXCR5-CCR6-CXCR3+CCR4-) cells map well to both Th1 and Temra cytotoxic^32, 33, 34^ clusters (**Figure 2**), coinciding with plasticity/lineage origin observations as discussed below.

Sorted Th17 (CCR10-CXCR5-CCR6+CXCR3-CCR4+) cells map to both Th17 and Th22 clusters, creating uncertainty that is successfully disentangled below.

Sorted Th1-17 (CCR10-CXCR5-CCR6+CXCR3+CCR4-) cells map to tight and linked zones within Th17 and Th1 clusters (**Figure 2, Figure S1B,C**). Of note, these zones do not coincide with any scRNA-Seq clusters obtained at any UMAP resolution (**Figure S6**), which warrants a deeper focused investigation.

Sorted Tfh (CCR10-CXCR5+) subset is found in two dissimilar scRNA-Seq clusters (designated as Tfh and Th1) differing in the expression of CXCR5 and CXCR3. This points at functional heterogeneity of Tfh subset^35^, and potential relations between Tfh and Th1 subsets^17^ which warrants a deeper investigation.

“Tnaive SOX4” cluster of Sakaguchi and colleagues^18^ corresponds to the cluster that we designated as “Naive RTE” (recent thymic emigrants), based on expression of CD31 (PECAM1)^36, 37^ and based on expression label transfer from umbilical cord blood CD4+ T cell scRNA-Seq^38^. “Tnaive act” cluster of Sakaguchi and colleagues^18^ corresponds to the cluster that we designated as “Naive”, based on the expression of BCL-2 involved in homeostatic proliferation of naive T cells^39^. “Tnaive” cluster of Sakaguchi and colleagues^18^ corresponds to the cluster that we designated as “Central memory 1”, based on the observed clonality (**Figure 3A**).

We also distinguished the “PD1^high^” cluster based on the expression of *PDCD1* and “cycling” cluster based on expression of *MKI67*

### Naive and effector IFN-induced clusters

Two of the peripheral Th scRNA-Seq clusters, designated as “Naive IFN-induced” and “Effector-memory IFN-induced” are clearly associated with type 1 interferon response. Effector-memory IFN-induced cluster, described earlier^42, 43^ is nearly absent in healthy donors’ blood but is well-detectable in COVID patients (**Figure 4A,B**), probably representing the typical behavior of Th cells in acute viral infection^43^. Notably, this cluster is almost undetectable in most critical COVID patients, which emphasizes the importance of type 1 interferon response in acute viral infection^44^. “Naive IFN-induced” cluster demonstrates similar behavior, and inversely correlates with the proportion of “Naive RTE” and “Naive” clusters, suggesting systemic and transient IFN-induced activation of conventional naive T cells in acute viral infection (**Figure 4A-C**). Similar observation of “Naive IFN-induced” cluster was recently reported on COVID and Systemic lupus erythematosus data^45^.

**Figure 4.**
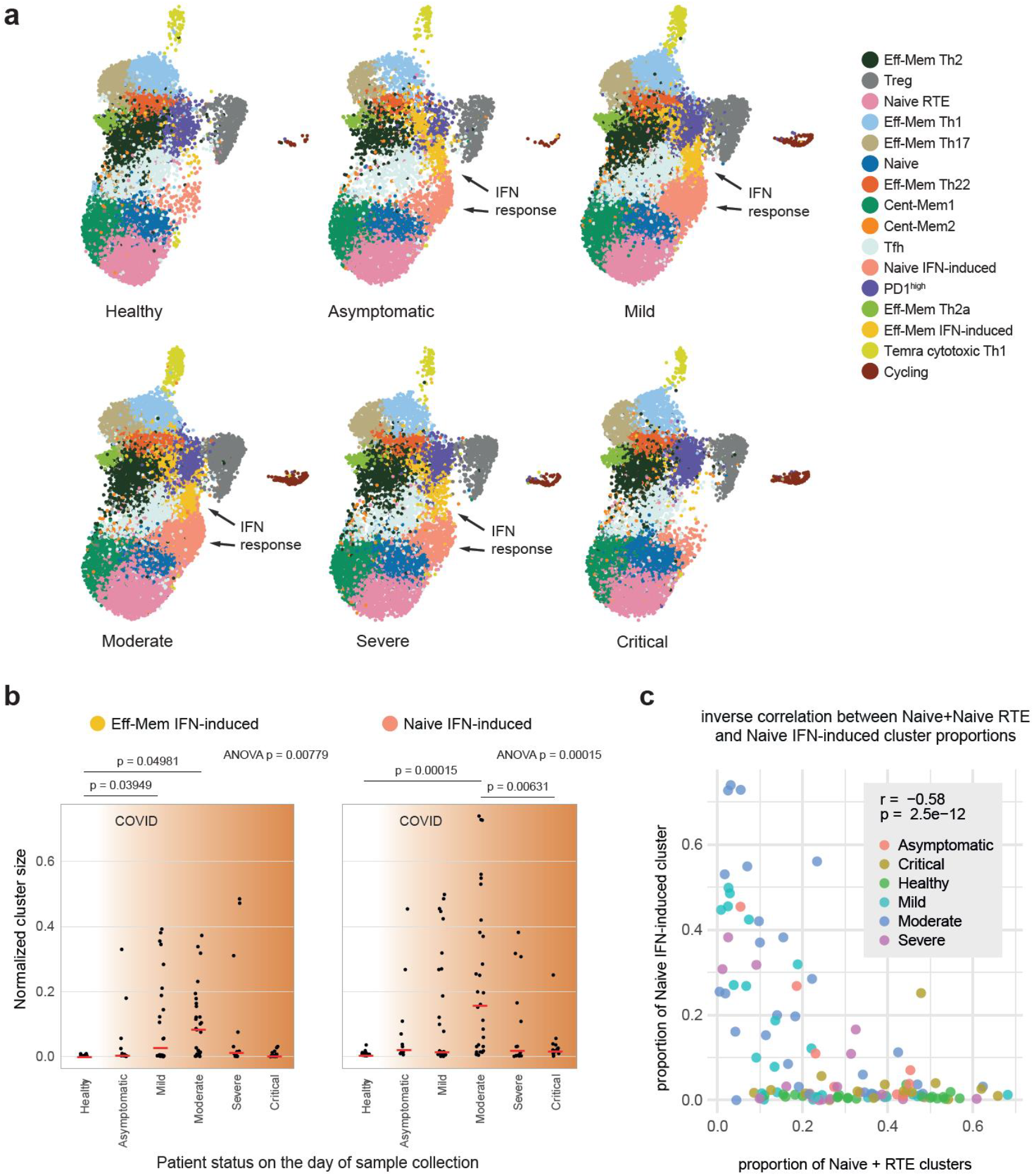
IFN response clusters in healthy donors and COVID patients. **a.** UMAP plots grouped by disease severity. “Eff-Mem IFN response” and “Naive IFN response” clusters are nearly absent in healthy individuals. **b,c**. Proportion of the samples occupied by “Eff-Mem IFN response” (b), or “Naive IFN response” (c) clusters (normalized cluster size). Medians are shown as red lines. p-values for ANOVA tests are shown on top; post-hoc pairwise analysis was performed with Tukey’s HSD test, and p-values < 0.05 are shown. **d**. The inverse correlation between proportions of Naive+Naive RTE versus Naive IFN-induced subsets within Th cells. Color shows patient status on the day of sample collection. r - Pearson correlation coefficient.

### Phenotypic versus intrinsic program plasticity of Th cells

To evaluate the relationships and cross-subset plasticity of T cell clones, we analyzed cluster stability at different clustering resolution levels (**Figure S6**) and clonal intersections between the clusters based on scTCR-Seq data (**Figure 3B**). We previously suggested notable plasticity between Th22/Th17, Th17/Th2, and Th2/Th2a subsets based on the corresponding intersections of sorted Th cell subset repertoires^28^. However, the new data on clonal overlaps and cluster stability at various resolutions in the current study prompts us to partially reconsider these interpretations.

Indeed, T cell clones sorted as classic Th17 subset (gated as NOT CD25^high^CD127^low^, CCR10-CXCR5- CCR6+CXCR3-CCR4+) are found in both Th17 and Th22 scRNA-Seq clusters. However, clonal overlap between these two clusters is low (**Figure 3B**), while the stability of both clusters is high (**Figure S6**). Furthermore, clones sorted as classic Th22 (gated as NOT CD25^high^CD127^low^, CXCR5-CCR10+) are almost exclusively found in Th22 scRNA-Seq cluster, indicating its self-standing nature. The independent origin of human Th22 clones was initially suggested^46^ and reported in mouse models^47, 48, 49^. Distinct TCR repertoire features also suggested the existence of clonally discrete Th22 subset^28^.

We interpret these observations as follows. The population classically sorted by phenotypic markers and described as Th17 actually represents the mixture of *bona fide* Th17 and Th22 cells (those of the latter that are stained as CCR10-negative). In contrast, the classically sorted CCR10+ Th22 subsets mostly coincide with the corresponding scRNA-Seq cluster, representing a relatively pure population of uniformly programmed *bona fide* Th22 T cells (**Supplementary Table 2**). To confirm this, we plotted those TCRβ clonotypes that were found as shared between sorted Th22 and Th17 subsets in Ref. 28, and these clonotypes were almost exclusively localized within the Th22 scRNA-Seq cluster. In contrast, sorted Th17 clonotypes that do not overlap with sorted Th22 predominantly mapped to Th17 scRNA-Seq cluster (**Figure S7**). That being said, some clonal overlap between Th2, Th2a, and Th22 scRNA-Seq clusters is observed (**Figure 3B**), thus leaving the room for long-term plasticity between these three subpopulations^49^.

Th1 and Temra cytotoxic Th1 clusters demonstrate notable clonal overlap (**Figure 3B**), indicating that the former could convert into the latter, as some of the recent works suggest^29, 30^. Correspondingly, the positioning of sorted Th1 TCRβ clonotypes to Th1 and Temra cytotoxic clusters may reflect terminal maturation of Th1 clones. Highest clonality was also observed for the cluster annotated as Temra cytotoxic Th1 subset, which is in line with the previous reports^33^ (**Figure 3A**).

In the proposed scRNA-Seq classification shown in **Figure 2A**, we give priority to the cells partitioning into stable scRNA-Seq clusters (**Figure S9**), while TCR-Track data are used to match classical surface phenotype-based sorted T cell subsets with those clusters.

### Programs of peripherally circulating Th clones are stable in time

Of note, T cell repertoires of the sorted subsets were obtained in 2018, while scRNA-Seq experiment was performed on samples derived in 2022. Despite the 4-year distance, TCR-Track mapped sorted clones to clearly defined positions, generally limited to one or two neighboring scRNA-Seq clusters (**Figure 2, Figure S1**), indicating long-term program stability of CD4^+^ T cell clones circulating in human peripheral blood.

Of course, the current behavior of T cell programs in tissues, in the context of the ongoing interaction with pathogens, microbiota, and environment can be associated with much more plastic behavior, forming a “continuum” where clear separation of Th programs may be problematic^26, 27, 50^. However, our results suggest that program imprinting may remain stable in most conditions, and when returning to the challenge-free circulation, the “resting” T cell memory clones return to their major initial imprint.

This concept is supported by generally low clonal overlap between scRNA-Seq clusters, with the exception of Th1 and Temra cytotoxic Th1 clusters, as well as natural intersection of “Cycling” Th cells with several differentiated Th clusters (**Figure 3B**).

## Discussion

The architecture of T cell memory essentially determines the entire pattern of our interaction with the antigens of the surrounding world, our microbiota, and our self-antigens^1, 2^. This architecture starts formation in the prenatal period^51, 52, 53, 54, 55^, is actively formed in the first years of life in the contact with pathogens, airborne and food antigens, and microbiota maturation^56, 57^ and then continues to be actively shaped by vaccinations and further contacts with infectious and non-infections challenges and antigens.

Clonal populations of T cells that are instructively primed by professional antigen-presenting cells^1, 2, 3^ make decisions about which reaction programs to choose to respond to each specific antigen. They remember these programs as memory clones, forming stable response patterns to familiar challenges, patterns of regulation and cross-regulation of immune responses to friends and foes.

Mistakes made in such decisions cost us dearly: they lead to autoimmune diseases, inefficient elimination of pathogens, chronic inflammation, cancer, and may essentially underlie the entire phenomenon of inflammaging. Presumably for this reason, some mammalian species are likely to avoid forming such a long-term clonal memory^11^.

In humans, however, the clonal memory of both CD8^+^ and CD4^+^ T lymphocytes can persist for years and decades^58, 59^. In the present work, TCR-Track allowed us to map Th subset repertoires on the very same T cell clones of the scRNA-Seq data obtained for the donor 4 years later, which emphasizes the stability of the huge number of clones accumulated over previous years^60^. We also show that these clones are mapped predominantly or even exclusively within their corresponding and independent scRNA-Seq clusters, highlighting the persistence of program decisions once made by each CD4^+^ T cell clone, as far as can be judged by peripheral blood analysis. Saying that, both in tissue plasticity^26, 27, 50^ and non-excluded opportunity for the multiple programs acquired by the progeny of a naive T cell that could be potentially revealed with deeper analysis^61^ stay in the game.

Repertoire-based TCR-Track annotation also turned out to be a powerful aid in matching classical subsets of lymphocytes sorted by surface markers with stable scRNA-Seq clusters that more comprehensively describe the diversity of functional lymphocyte programs. This effort structures our understanding of functional diversity of Th cells, critical for further progress in cancer and autoimmunity immunotherapy and vaccine development.

CITE-seq can also be used to phenotype cell populations on a single-cell level based on surface markers, having an advantage of simultaneously measuring hundreds of those. Our work shows that TCR-Track may outperform CITE-Seq in terms of resolution capacity. At the same time, we should indicate that this may be attributed to a technical instability in multiplex CITE-Seq, where successful CITE-Seq experiment may yield results comparable to TCR-Track. Additionally, some of the CITE-Seq staining antibodies may affect cell signaling and transcriptomic profiles, introducing bias in the scRNA-Seq landscape^62^. In contrast, in the TCR-Track pipeline, cells stained with surface antibodies for sorting and bulk TCR-Seq are obtained and analyzed independently of the scRNA-Seq/scTCR-Seq experiment.

We anticipate that in the future works, TCR-Track can be used to match and classify: 1) CD4^+^ T cell populations in the peripheral blood, lymph nodes, tertiary lymphoid structures at the sites of chronic inflammation and tumor environment, other tissues in health and disease; 2) diverse follicular helper T cells^35^; 3) known and unknown types of invariant and semi-invariant T cells, such as iNKT, NKT, MAIT, and CAIT^63, 64, 65^; 4) gamma delta T cells^66^; 5) central memory and stem cell memory T cells that would probably require deeper scRNA-Seq coverage due to their relatively lower clonality; 6) CD8^+^ T cell subsets^67^; 7) B cell functional subsets^68^.

TCR-Track approach has certain limitations that must be taken into account. Firstly, similar to CITE-Seq, the method relies solely on surface markers to define target cell subpopulations. Secondly, the method is applicable only to T- and B-cells with specific clonal receptors acting as living barcodes. Thirdly, TCR- Track is based on lymphocyte clonality, where we need to capture the same clone in sorting and scRNA-seq experiments. Therefore, TCR-Track based annotation of e.g. central memory T cell subsets or peripheral blood Tfh cells, which are almost as diverse as naive T cells^28, 35^, may require substantially deeper scRNA-Seq and bulk TCR profiling. For the naive T cell subsets, TCR-Track application would be probably limited to innate-like, relatively clonal naive T cell subpopulations of fetal origin^53^.

Summarizing, our work:

1. Offers TCR-Track as an approach for classification of phenotypically defined T- and B-lymphocyte subsets, their exact positioning within scRNA-Seq landscape and time-tracking.
2. Clarifies correspondence between the well-studied human peripheral blood Th subsets and scRNA-Seq clusters.
3. Proposes more accurate, clonally informed CD4^+^ T cell classification within scRNA-Seq landscape, delineating positioning of Th2, Th2a, Th22, and Tfh clusters, and refining several other details. An integrated scRNA-Seq reference dataset of peripheral Th lymphocytes is provided.
4. Shows long-term program stability and low intrinsic plasticity of Th memory clones circulating in peripheral blood.
5. Shows high clonal overlap suggesting that cytotoxic CD4+ T cells differentiate from Th1 clones.
6. Shows that Th17 and Th22 represent clonally independent subsets. Sorted Th17 subset represents a mixture of *bona fide* Th17 and CCR10^low^ Th22 cells.
7. Shows that SARS-CoV-2 infection is associated with transient IFN type 1 activation of naive T cells, the phenomenon which warrants a deeper investigation.
8. Shows that efficient response to SARS-CoV-2 infection is associated with appearance of a prominent effector IFN-induced Th cells, while critical COVID is associated with low presence of the IFN-induced subset.

More generally, we hope that this work advances the study of the role of programmed populations of memory T cells to a new level, making it possible to clearly distinguish the functional nature of each immune response, as well as to investigate the plasticity between T cell subsets. This level of understanding serves as a necessary steppingstone to the rational development of better immunotherapeutic approaches in oncology and autoimmunity, and in vaccine development, where the chosen T cell programs fundamentally determine the type of immune response and are crucial to the outcome.

## Acknowledgements

We thank Vadim Karnaukhov, Denis Syrko, Viktor Kotlyar, Ivan Pozdniakov, and Dmitry Bolotin for assistance with data analysis, and Maria Vakhitova, Tatiana Grigorova, Ilgar Mamedov, and the NGS Laboratory (IKMB, Kiel University) for assistance with sequencing, which was also supported by the DFG Research Infrastructure NGS_CC (project 407495230) as part of the Next Generation Sequencing Competence Network (project 423957469). The work was supported by the Ministry of Science and Higher Education (grant no. 075-15-2019-1789).

## Author contributions

Study conception: D.M.C. Provided peripheral blood mononuclear cells (PBMCs) and enrolled donors: K.L., B.E.M., A.Y.K., D.A.P. Isolated PBMCs, performed T cell sorting, bulk TCR-Seq, scRNA-Seq and CITE-Seq experiments: V.V.K, K.L, I.A.S., D.B.S., B.E.M., A.S.F., P.S., A.Y.K., K.L.M. Sequencing: V.V.K., K.A.Y., A.F. Data analysis: D.K.L., V.V.K., O.N.S., P.S., A.Y.K., K.A.B. Manuscript writing: D.K.L., V.V.K., D.M.C. Project coordination: V.V.K., A.Y.K., B.O.V., A.F., D.A.P., D.M.C.

## Declaration of interests

The authors report no competing or conflicting interests.

## Data and code availability

Raw sequencing data is available in NCBI Sequence Read Archive (BioProject: PRJNA995237). A processed Seurat object with the Th reference (“full_reference_return_model.rds”) is available for download at https://figshare.com/projects/T_helper_subsets_Kriukova_et_al_/173466.

Code for figures is available at https://github.com/kriukovav/Thelpers.

## Method details

### scRNA-Seq and scTCR-Seq library preparation and sequencing

Fresh PBMCs from D01, D04, D05 were stained with anti-CCR7–PE-Cy7 (clone 3D12, BD Biosciences), anti-CD3–APC-Fire750 (clone SK7, BioLegend), anti-CD4–PE-Cy5.5 (clone S3.5, Thermo Fisher Scientific), anti-CD14–V500 (clone M5E2, BD Biosciences), anti-CD19–V500 (clone HIB19, BD Biosciences), anti-CD45RA–PE-Cy5 (clone HI100, BioLegend), and LIVE/DEAD Fixable Aqua (Thermo Fisher Scientific). Viable effector/memory CD4^+^ T cells gated as CD3^+^ CD4^+^ CD14^−^ CD19^−^ after excluding CCR7^+^ CD45RA^+^ events were sorted in two replicates for D01 and without replicates for D04 and D05 using a custom-modified FACSAria II (BD Biosciences) and loaded onto a Chromium Controller (10x Genomics). Samples were prepared using a Chromium Next GEM Single Cell 5’ Reagent Kit v2 (10x Genomics). Pooled samples were sequenced with a coverage of 100,000 reads per input cell for scRNA-Seq and 25,000 reads per input cell for scTCR-Seq on a NovaSeq 6000 System with an S4 Flow Cell (Illumina).

### scRNA-Seq, scTCR-Seq and CITE-seq data analysis

Raw scRNA-Seq and scTCR-seq fastq files were processed using the *count* and *vdj* pipelines in cellranger (v6.1.2). For scTCRs, only productive in-frame CDR3- and V-J-spanning contigs without the stop codons in the V-J region were selected, and the most abundant TCR chain was used for cells with two relevant transcripts (TRA or TRB). Filtered gene expression matrices were uploaded into the Seurat R package (v4.2.0)^69^. The cells containing >10000 UMIs and >10% mitochondrial reads were removed. The data from each donor was paired with the corresponding scTCR data, log normalized with *NormalizeData* function and clustered with the Louvain algorithm. Publicly available processed multimodal scRNA-seq/scTCR-seq/CITE-seq data from PBMCs were downloaded from the ArrayExpress database under ascension number E-MTAB-10026. This data was converted to Seurat object, applying the same filtering criteria as used for the newly generated data. CD4^+^ T cell clusters were selected based on the average cluster expression of *CD4* and *CD3E* and the cluster annotations provided by the authors. The public dataset was then split by sample origin, log normalized and clustered with the Louvain algorithm based on sets of highly variable genes calculated individually for each batch with Seurat functions. CITE-Seq data, provided as ADT counts for 192 features in processed data, were normalized separately for each batch using the centered log ratio (CLR) method as implemented in Seurat and used for further analysis. In addition to filtering steps originally performed by the authors, at this stage we removed outlier clusters with both low UMI counts and high percentage of mitochondrial genes, as well as clusters expressing *CD8A, CD8B* from all individual datasets. Differential expression was calculated using *FindAllMarkers* function individually on each dataset. Clusters that either highly expressed or contained more several of B cell, dendritic cell, macrophage and other non T cell markers in the differential expression as per single-cell RNA section of Human Protein Atlas https://www.proteinatlas.org/humanproteome/tissue were removed from the analysis, clusters that expressed the markers of NK cells without the expression of CD3/TCR were also removed. The stress score was calculated for each cell using *AddModuleScore* function in Seurat, it included the genes (BTG1, BTG2, DDX5, DNAJA1, DUSP1, EEF1A1, HSPA8, JUN, JUNB, JUND, KAP, KLF6, PNRC1) upregulated as a consequence of the dissociation procedure^70^.

The paired scRNA/scTCR-seq dataset generated in this study contains 23257 cell barcodes, of which approximately 90% contains scTCR-seq data in each of the 3 donors. The total cell counts for the donors are 4723, 9430, and 9104. For public data, out of 124420 cell barcodes from the 119 donors selected for the analysis approximately 80% contained scTCR-seq data. In public data, 20% of cells came from 21 healthy donors, 77% of cells came from 93 donors with COVID-19 infection across several disease severity groups: asymptomatic (9400 cells), mild (29807 cells), moderate (24774 cells), severe (14303 cells) and critical (17031 cells), as well as 3% of cells from 5 donors with non-covid infection.

Integration of scRNA-seq data was carried out using the Seurat reference-based reciprocal PCA protocol with default parameters, with the largest 3’ (Newcastle public data) and 5’ (D05) datasets chosen as references to account for differences in methodology^69^. The percentage of mitochondrial genes and the stress score were regressed out of the integrated dataset as implemented in *ScaleData* function. The number of dimensions used for running the UMAP and Louvain clustering algorithm was 25 based on the *ElbowPlot* Seurat function. All TCR and IG genes were removed from variable features used in the PCA and from the anchor features used for integration. To identify cluster marker genes, *FindMarkers* and *FindAllMarkers* function were used on the scRNA-seq data slot.

For the label transfer from UCB data, we used processed T cell scRNA-seq data from figshare (https://figshare.com/projects/Single-cell_mapping_of_progressive_fetal-to-adult_transition_in_human_naive_T_cells/76143), and gated CD4 T cell clusters from umbilical cord blood as described previously. We projected the cluster identities of the integrated data onto this dataset via the Seurat data transfer method (*FindTransferAnchors* and *TransferData* functions). The distribution of predicted IDs with the confidence score > 0.5 was used to support the annotation of cluster 2 as “Naive RTE” (data not shown).

*In silico* CITE-seq gating was performed sequentially, following the scheme in **Figure S5** and visualised using the R packages ‘ggplot2’ (v3.4.4) and ‘ggpointdensity’ (v0.1.0). CCR10 marker expression was assessed using scRNA-Seq since CCR10 was not included in the CITE-Seq panel. To quantitatively evaluate the accuracy of TCR-track and CITE-seq cluster mapping, Normalized Shannon-Wiener index 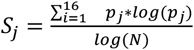 was calculated, where *j* is the gated subset as in **Figure S5** (e.g. Tfh, Th1, Th1-17), *i* is the scRNA-seq cluster (e.g. 0, 1, 2), p_j_ is the proportion of gated subset *j* in cluster *i*, and N is the total number of clusters *i* with non-zero hits for the gated subset *j*. It measures the evenness of cell distribution across clusters for each gated subset. The gates for CITE-seq gating were chosen to minimize the resulting normalized Shannon-Wiener index for each gated subset. For the result evaluation, the index was separately calculated within three donors in TCR-track data, and within the five disease severity groups in public CITE-seq data (donors were grouped to make the number of cells comparable in each of the analyses). Only the gated subsets with more than 7 cells were included in the analysis.

### Mapping of the FACS-sorted Th subsets’ clonotypes

From each bulk TCRβ repertoire (Th1, Th2, Th17, Th1-17, Treg, Th2a, Tfh, Th22) from Kasatskaya et. al. (Ref. 28) the top-500 nucleotide clonotypes were selected, to make analysis uniform, and also to exclude even minor cross-contaminations that could happen during cell sorting or sequencing. In cases when replicates were available for the bulk TCR repertoires, only one of the pairs was used in the analysis. Then, for each single cell from D01, D04, and D05 the respective TCRβ clonotype frequencies from top-500 bulk TCR-Seq Th datasets were added to the Seurat object metadata. These frequencies were used as the transparency encoding variables (aesthetic “alpha” in ggplot2) on the **Figure 2C, Figures S1B** to emphasize the highly abundant within the respective Th subsets clonotypes. In the **Figures S1C-D, S10** all shown cells have an identical value of transparency.

### Identification of clonotypes overlapping between sorted Th17 and Th22 subset repertoires

The overlapping TCRβ nucleotide clonotypes were identified between the sets of top-2,000 TCR clonotypes from the sorted Th17 and Th22 cell subsets^28^. Then a standard deviation was calculated for the frequencies of each overlapping clonotype in the Th17 and the Th22 subsets from the same donor. We selected only those overlapping clonotypes, for which this standard deviation was lower than 0.001. This way from the list of overlapping clonotypes we excluded those which were highly abundant in the Th17 subset and low abundant, but still present, in the Th22 subset, or *vice versa*.

### Ethics statement

Ethical approval was granted by the Cardiff University School of Medicine Research Ethics Committee (16/55).

## Notes

### Competing Interest Statement

The authors have declared no competing interest.

### Summary of Updates

For the sake of clarity we keep only a part of the original submission in this version of the manuscript. The part of the original work related to CultivAToRR will be published separately.

