## Supplementary Figures and Supplementary Table2 for "Repertoire-based mapping and time-tracking of helper T cell subsets in scRNA-Seq"

**Supplementary Table 2**. Correspondence between classically sorted peripheral blood Th subsets and stable scRNA-Seq clusters, according to Sort-Seq.

| **Classic name** | **FACS-sorted, gated as CD4+, NOT CCR7+CD45RA+ (effector-memory CD4+ T cells)** | **Correspondence to stable scRNA-Seq clusters (*bona fide* Th programs)** |
|---|---|---|
| Th1 | NOT $CD25^{high}CD127^{low}$, CCR10-CXCR5-CCR6-CXCR3+CCR4- | 5:1 mixture of Temra cytotoxic Th1 and Th1 clusters |
| Th1-17 | NOT $CD25^{high}CD127^{low}$, CCR10-CXCR5-CCR6+CXCR3+CCR4- | Independent subset located within linked and tight zones of Th1 and Th17 scRNA-Seq clusters, requires deeper investigation |
| Th17 | NOT $CD25^{high}CD127^{low}$, CCR10-CXCR5-CCR6+CXCR3-CCR4+ | 3:2:2 mixture of Th22, Th17 and Th2 clusters |
| Th22 | NOT $CD25^{high}CD127^{low}$, CCR10+CXCR5- | 7:2 mixture of Th22 and Th2 clusters |
| Th2 | NOT $CD25^{high}CD127^{low}$, CCR10-CXCR5-CCR6-CXCR3-CCR4+CRTh2- | 6:1 mixture of Th2 and central memory clusters |
| Th2a | NOT $CD25^{high}CD127^{low}$, CCR10-CXCR5-CCR6-CXCR3-CCR4+CRTh2+ | 3:1 mixture of Th2a and Th2 clusters |
| Treg | $CD25^{high}CD127^{low}$ | Treg clusters, localized in the zone of “effector Tregs” according to classification by Sakaguchi and colleagues[1] |
| Tfh | NOT $CD25^{high}CD127^{low}$, CCR10-CXCR5+ | Tfh, Th1 (presumably Tfh1), requires deeper investigation |

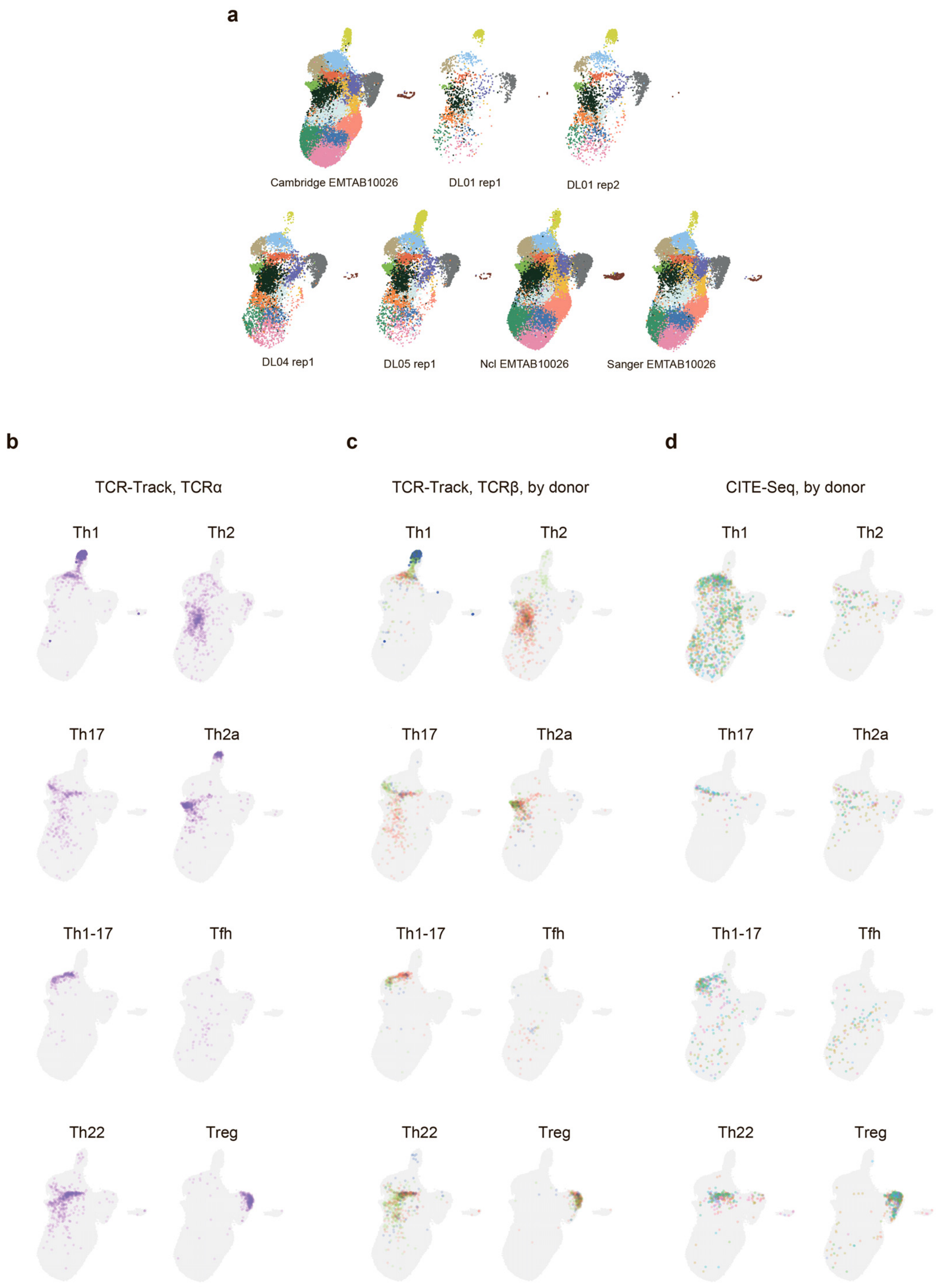


**Supplementary Figure 1. Building and characterization of the reference dataset. a.** UMAP plots of integrated dataset split by the origin of samples, colored by the assigned cluster. **b**. UMAP plots showing the localization of Sort-Seq-defined TCRα clonotypes. **c**. UMAP plots showing the localization of Sort-Seq-defined TCRβ clonotypes, colored by donor (n = 3). **d**. UMAP plots showing the localization of CITE-Seq-defined cells, gated *in silico* as shown on **Supplementary Fig. 5,** colored by donor (n = 119).

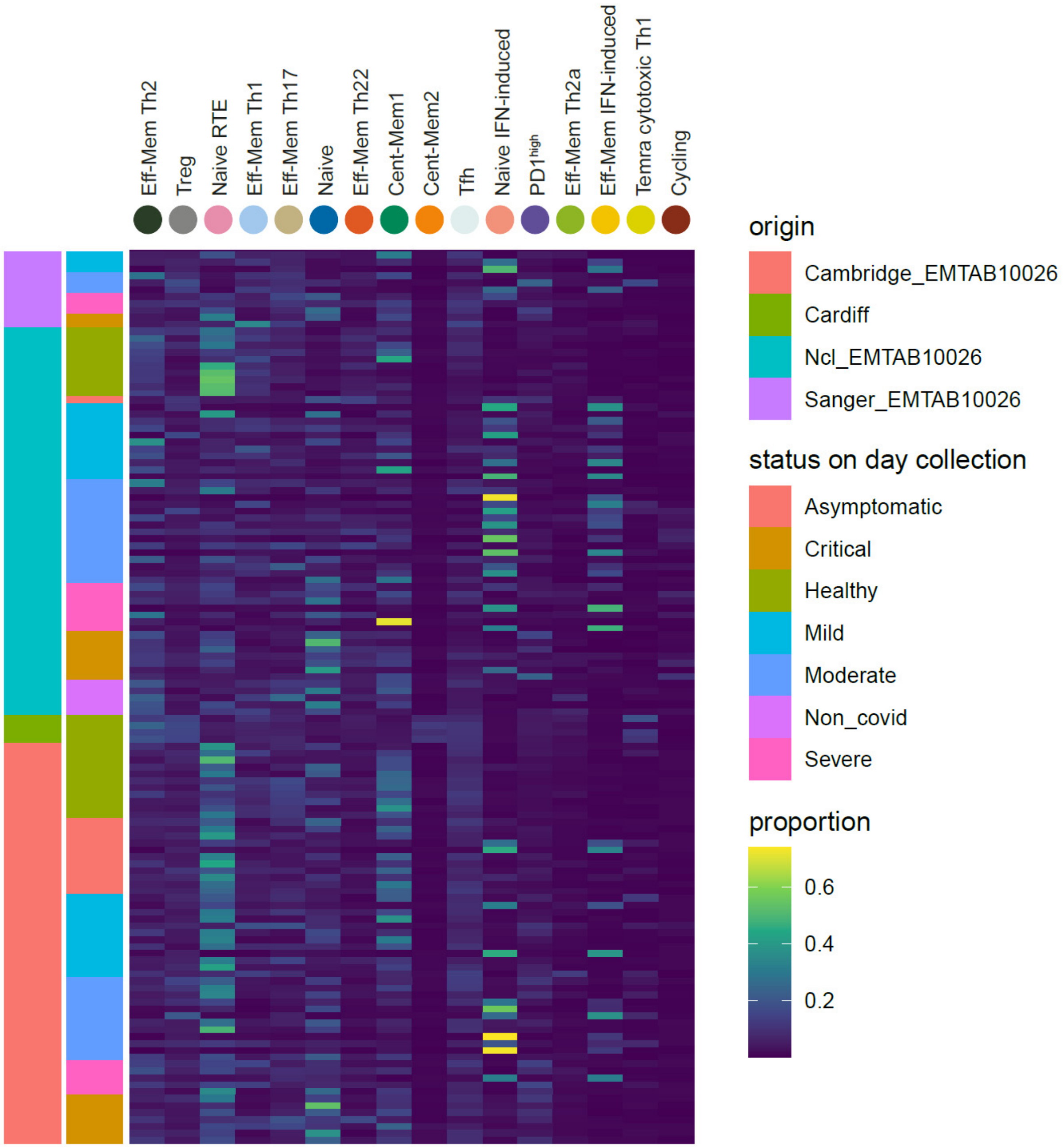


**Supplementary Fig. 2. scRNA-Seq clusters distribution across studies and COVID disease status.**

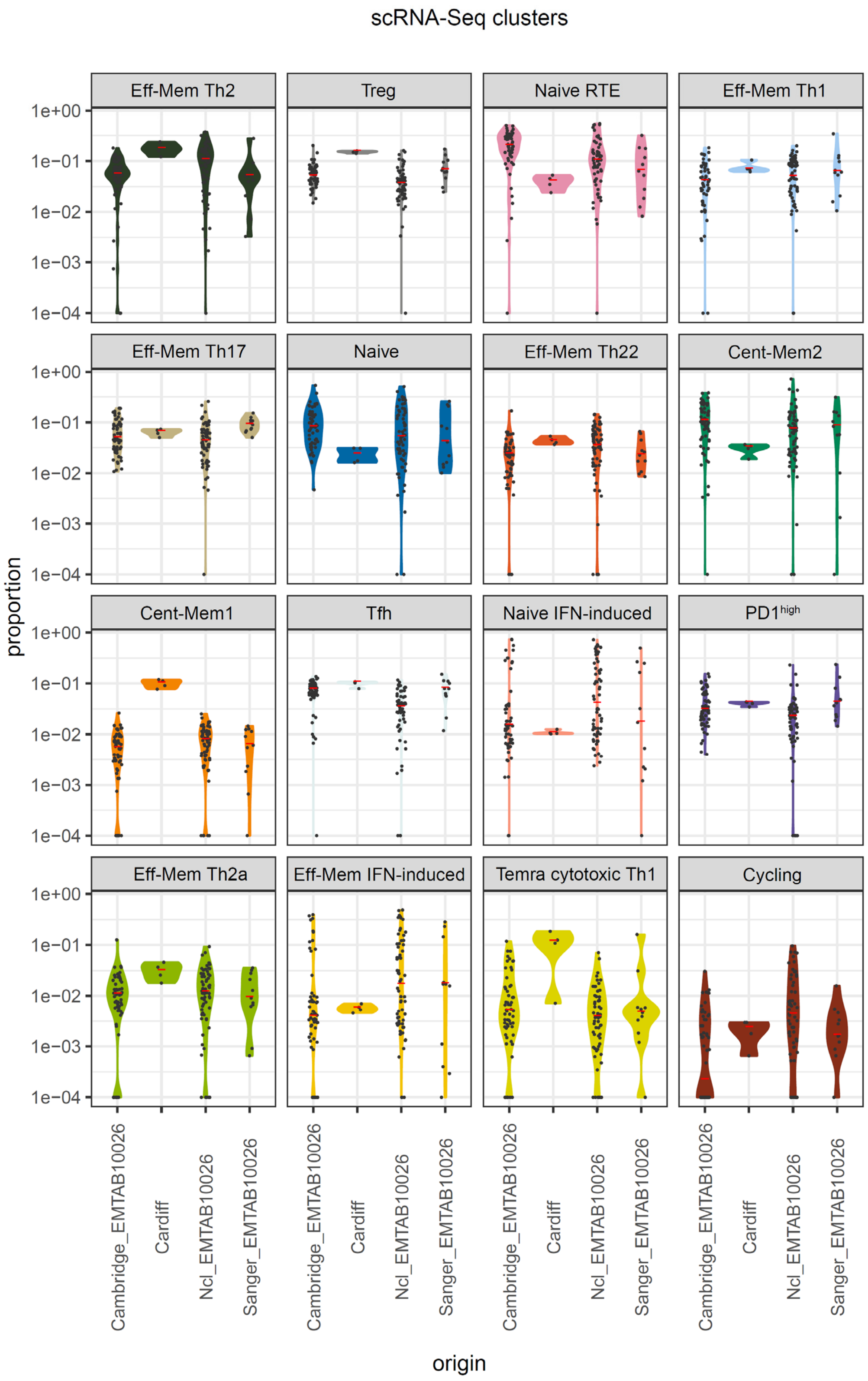


**Supplementary Fig. 3. scRNA-Seq clusters distribution across studies.**

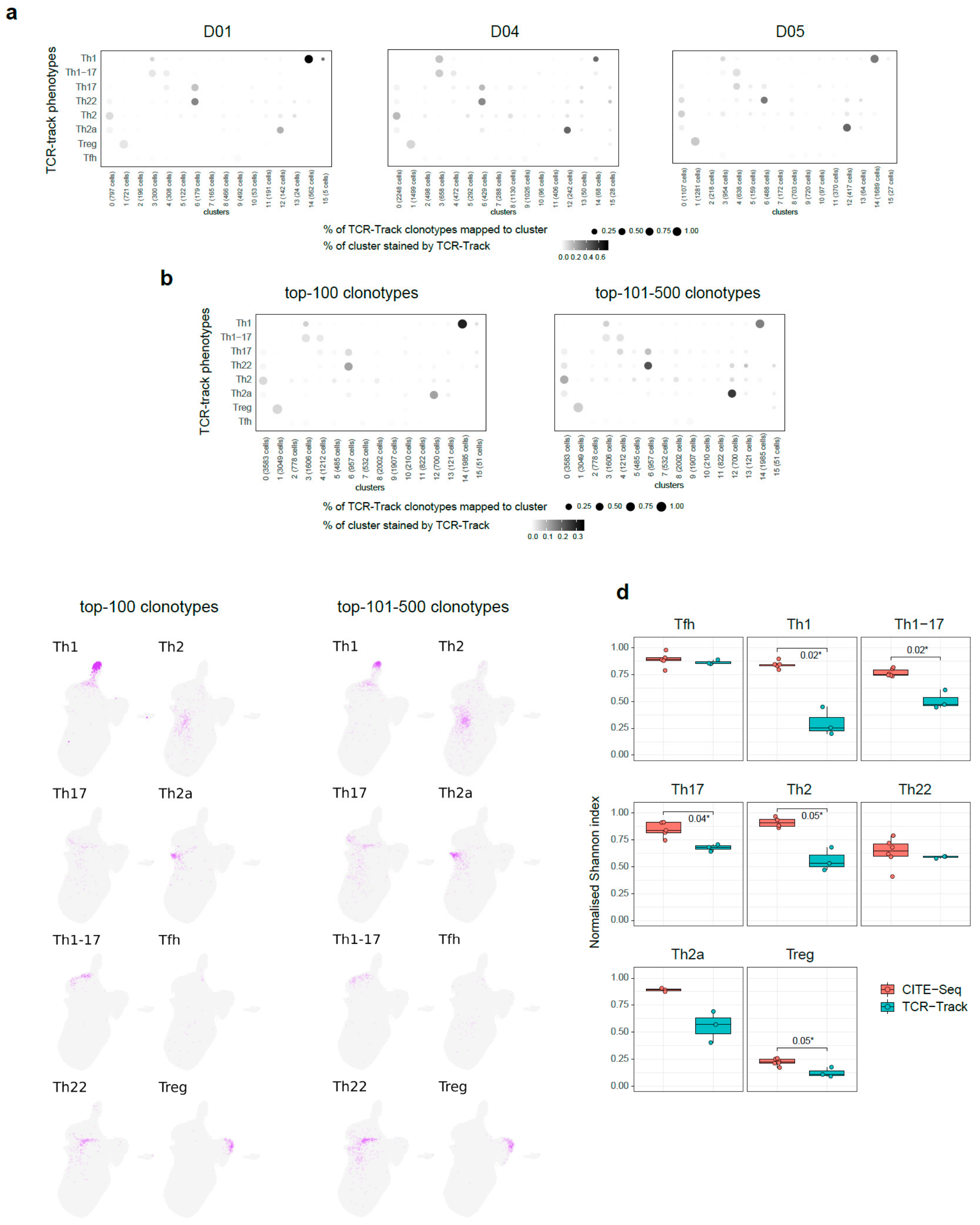


**Supplementary Fig. 4. TCR-Track reproducibility and clonality dependence. a.** Dot plots summarizing the clonal positioning of sorted Th subsets (TCR-Track) within the scRNA-Seq clusters, shown separately for each donor. **b.** Dot plots summarizing the clonal positioning of top-100 and top101-500 largest clonotypes from the sorted Th subsets (TCR-Track) within the scRNA-Seq clusters. **c.** UMAP plots showing the clonal positioning of top-100 and top101-500 largest clonotypes from the sorted Th subsets (TCR-Track) within the scRNA-Seq clusters. TCRβ clonotypes were used to define Th subsets in TCR-Track method. The color intensity in TCR-Track is proportional to the clonal frequencies in the original sorted Th bulk TCRβ repertoires. **d.** Relative accuracy of subsets mapping to the specific scRNA-Seq clusters measured as Normalized Shannon index that reflects unevenness of cell distribution across clusters (Wilcoxon rank sum test, see Methods). The lower the value, the more focused is the mapping.

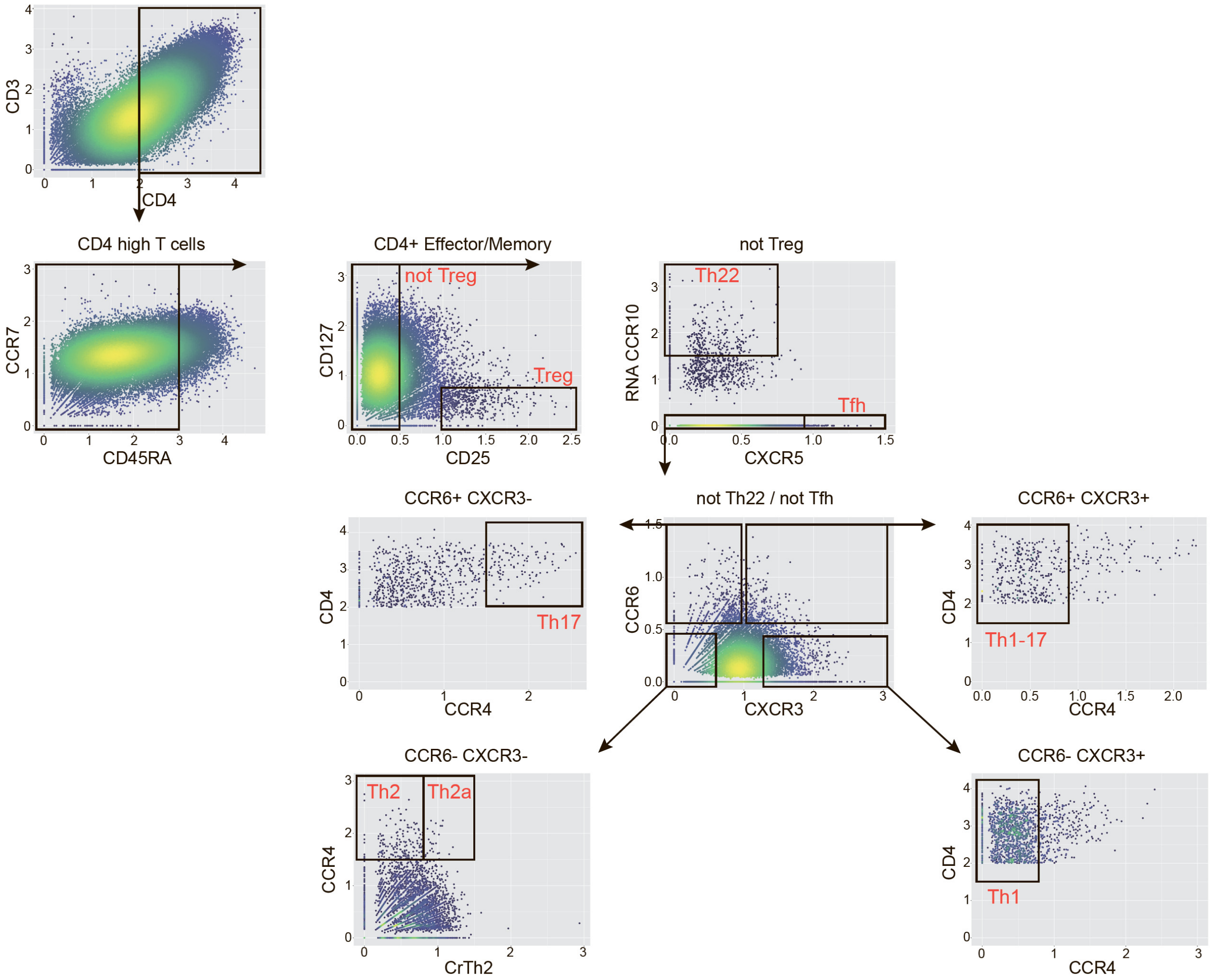


**Supplementary Figure 5. *In silico* flow-cytometry-like gating strategy.** *In silico* gating with CITE-seq "flow-cytometry-like" biplots corresponding to FACS gating scheme. Only samples with CITE-seq data are included. CCR10 expression is measured by scRNA-Seq due to the absence of CCR10 in the CITE-Seq panel. Color corresponds to the number of neighbor points.

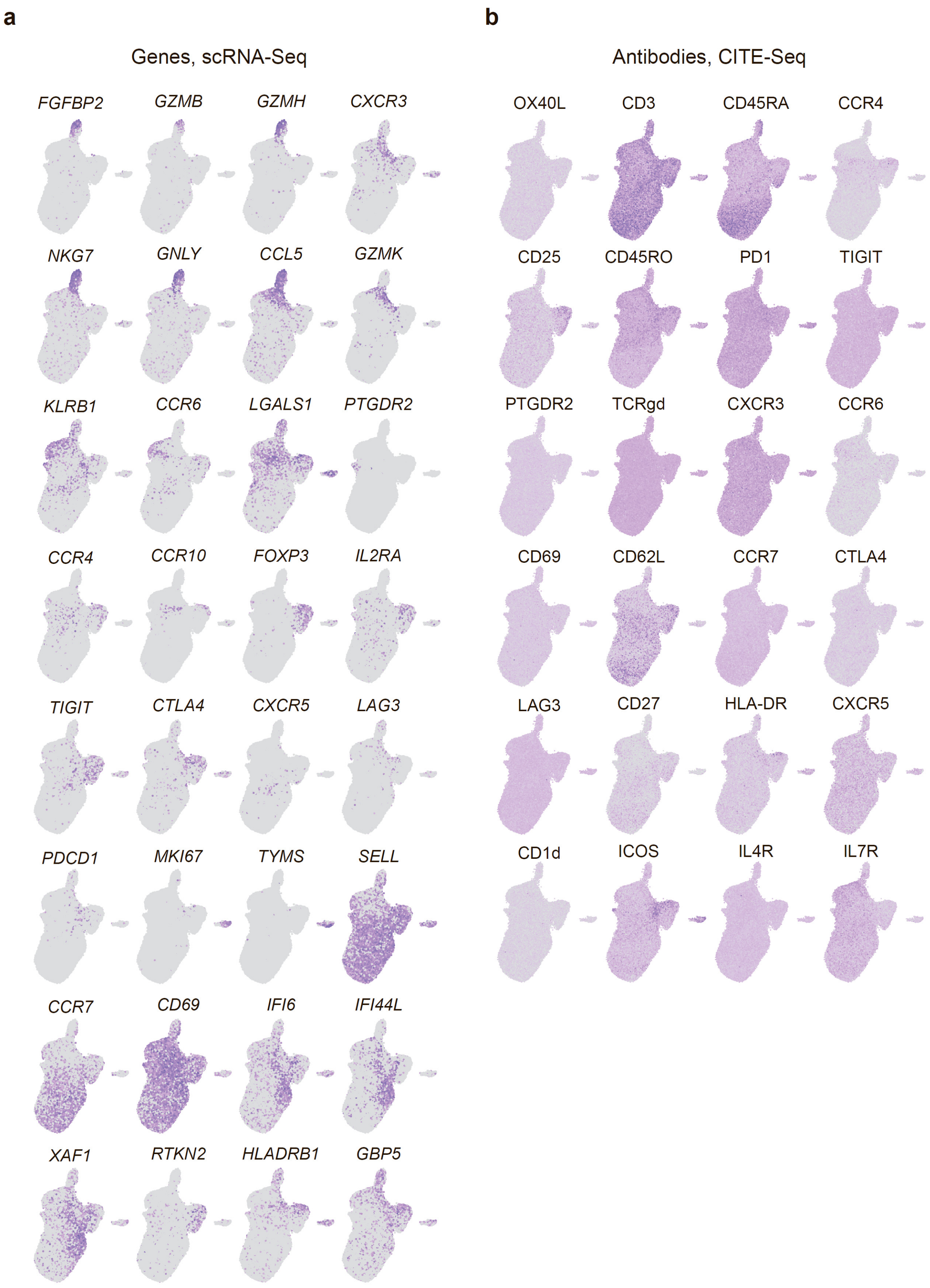


**Supplementary Figure 6. Marker gene and protein expression in the reference dataset. a.** Marker genes, scRNA-Seq. **b.** Marker proteins, CITE-Seq.

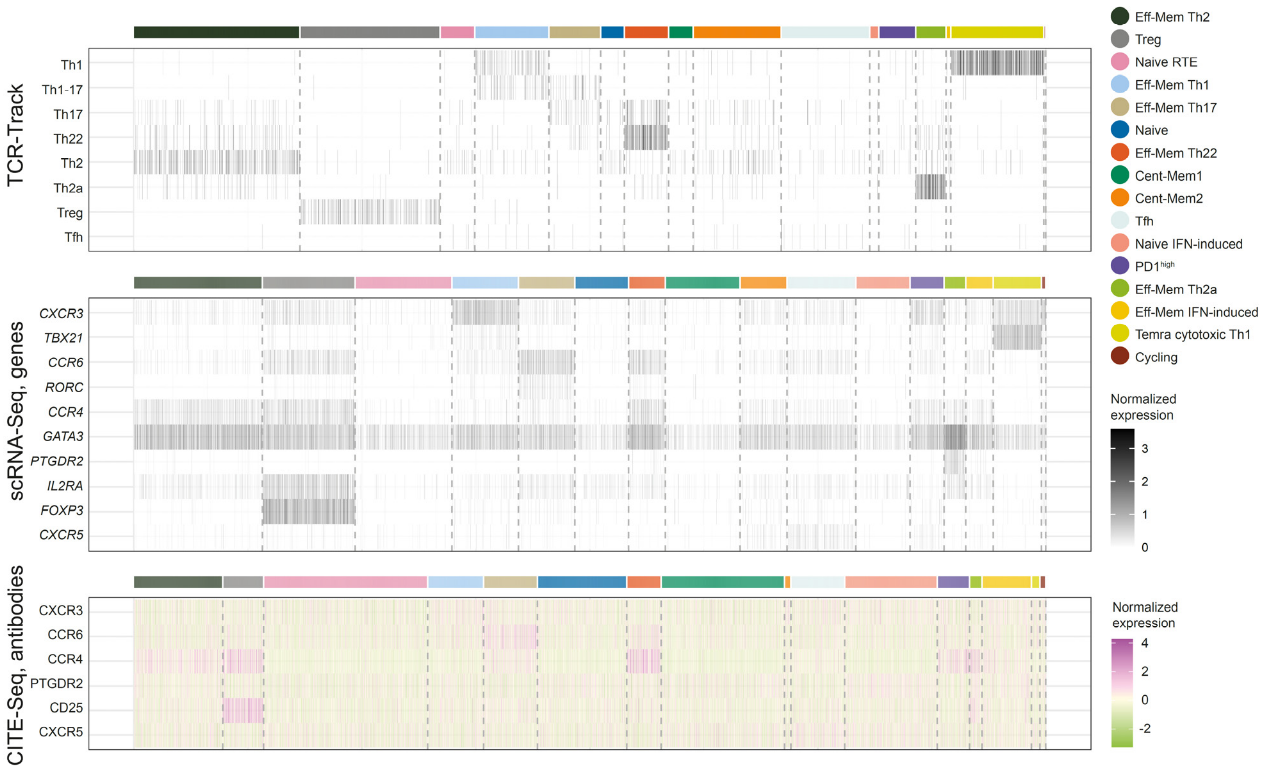


**Supplementary Fig. 7. Characteristic genes and proteins.** Heatmap summarizing the positioning of TCR-Track clonotype-determined subsets, scRNA-Seq expression of characteristic genes, and CITE-Seq signal for the corresponding surface proteins. For normalization, equal numbers of 10,000 scRNA-Seq cells were randomly selected from TCR-Track and CITE-Seq experimental datasets. Each tile of the heatmap represents one cell. The color intensity in the scRNA-Seq plot visualizes the gene expression in a cell (white color depicts zero expression). Protein expression measured by CITE-Seq was Z-scored, and all values exceeding 99.5 percentile were trimmed.

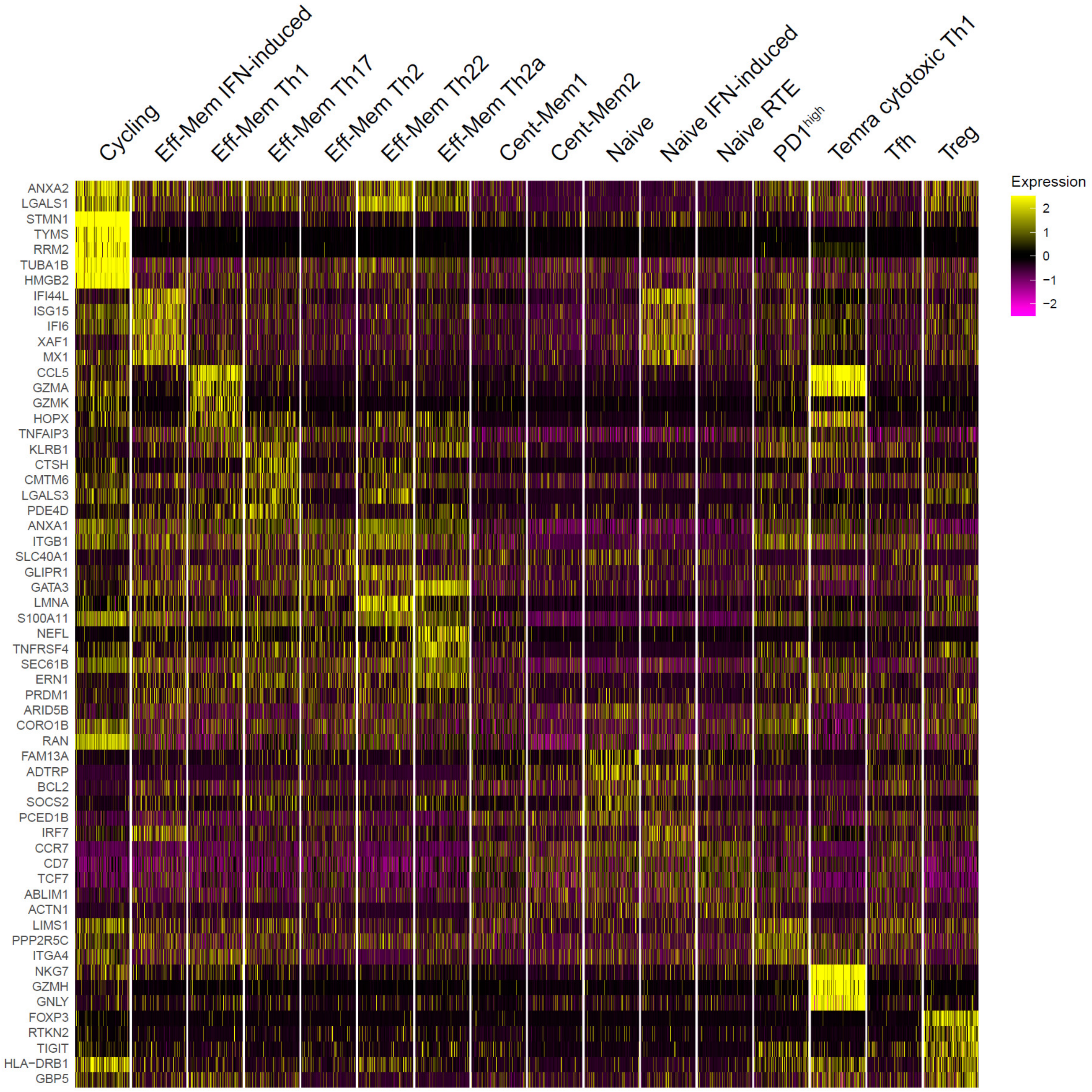


**Supplementary Fig. 8. Top genes that distinguish peripheral Th scRNA-Seq clusters.**

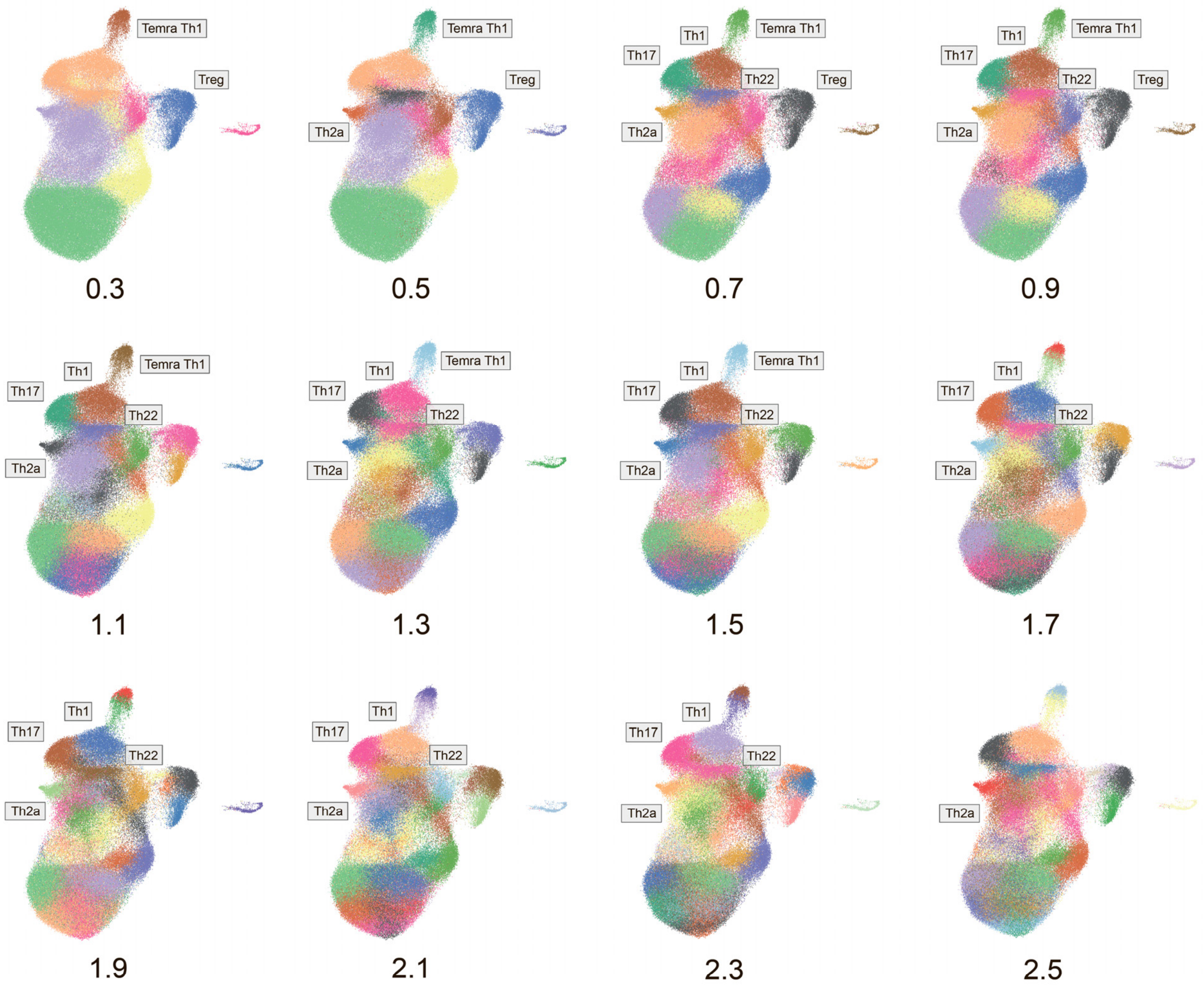


**Supplementary Fig. 9. Stability of Th programs studied through scRNA-Seq data.** UMAP plots built with different clustering resolutions. Th1, Th17, and Th22 clusters are stable and conserved across various resolutions.

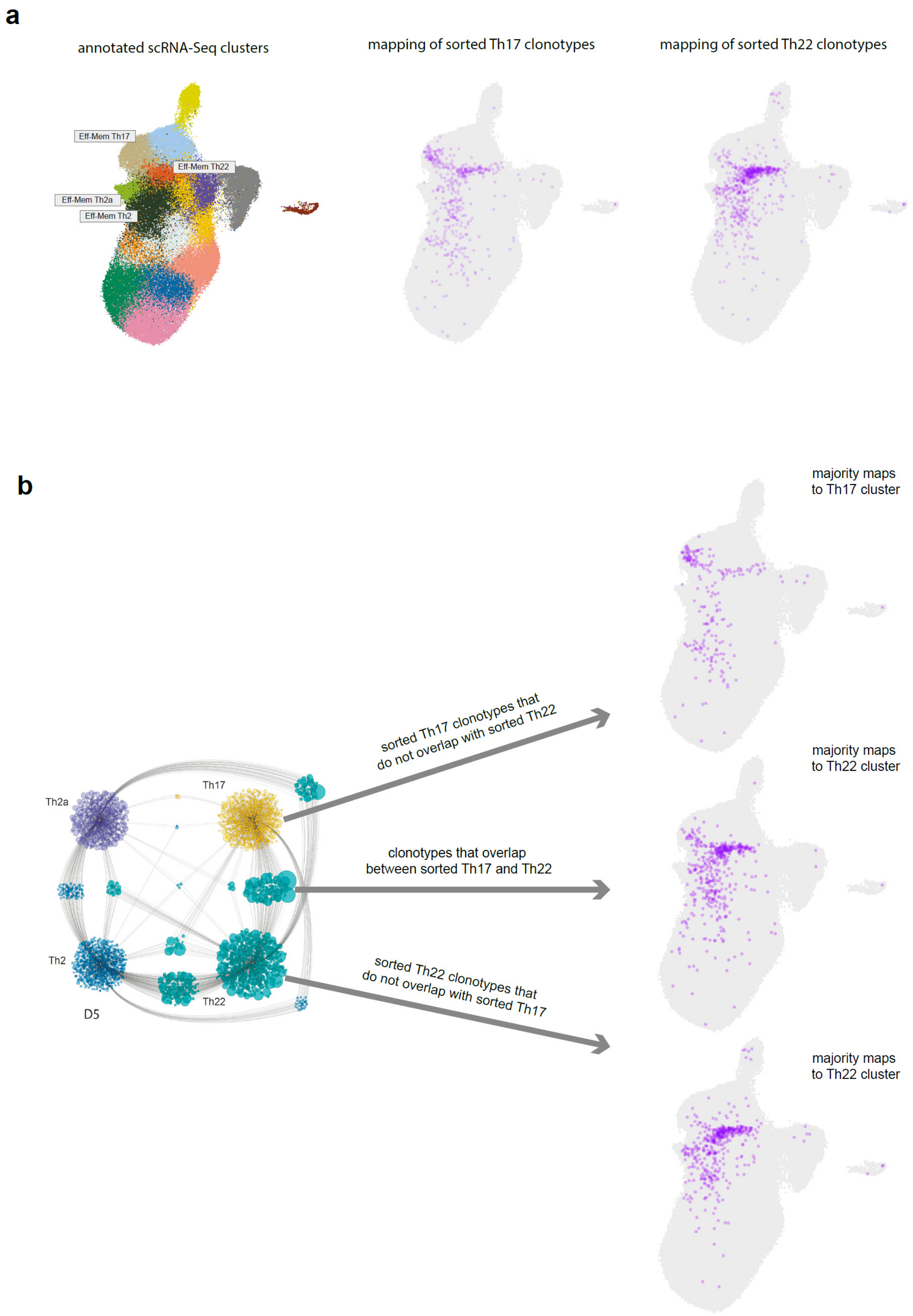


**Supplementary Fig. 10. Positioning of the Th17/Th22 shared clonotypes. a.** Annotated scRNA-Seq clusters and TCR-Track positioning of sorted Th17 and Th22 subsets. **b.** UMAP positioning of sorted Th17 clonotypes that do not overlap with sorted Th22, sorted Th22 clonotypes that do not overlap with sorted Th17, and overlapping clonotypes. Cytoscape network plot adapted from Ref. [2].
